# Multi-method brain imaging reveals impaired representations as well as altered connectivity in adults with dyscalculia

**DOI:** 10.1101/162115

**Authors:** Jessica Bulthé, Jellina Prinsen, Jolijn Vanderauwera, Stefanie Duyck, Nicky Daniels, Céline R. Gillebert, Dante Mantini, Hans P. Op de Beeck, Bert De Smedt

## Abstract

Two hypotheses have been proposed about the etiology of neurodevelopmental disorders: representation impairments versus disrupted access to representations. We implemented a multi-method brain imaging approach to directly compare the representation vs. access hypotheses in dyscalculia, a highly prevalent but understudied neurodevelopmental disorder in learning to calculate. We combined several magnetic resonance imaging methods and analyses, including multivariate analyses, functional and structural connectivity, and voxel-based morphometry analysis, in a sample of 24 adults with dyscalculia and 24 carefully matched controls. Results showed a clear deficit in the non-symbolic magnitude representations in parietal, temporal, and frontal regions in dyscalculia. We also observed hyper-connectivity in visual brain regions and increased grey matter volume in the default mode network in adults with dyscalculia. Hence, dyscalculia is related to a combination of diverse neural markers which are altogether distributed across a substantial portion of cerebral cortex, supporting a multifactorial model of this neurodevelopmental disorder.

## 1. Introduction

Hypotheses about the neural basis of neurodevelopmental disorders fall into two categories: dysfunction in specific brain regions and cognitive representations, versus disrupted connectivity and impaired access to these representations. This distinction runs through the history of neuroscience, starting when the localized function approach in language disorders was appended by the concept of disconnection syndromes as introduced by Wernicke (Spreen et al., 1995).

Recently, a multi-method brain imaging approach was used in adults with developmental dyslexia to directly compare the representation versus access hypotheses by combining measures of the quality of neural representations with measures of connectivity (Boets et al., 2013). This study revealed that dyslexia in adults is associated with disrupted connectivity without dysfunctional representations (Boets et al., 2013). Here, we test whether this can be extrapolated to other neurodevelopmental disorders, specifically to developmental dyscalculia, a disorder in learning to calculate.

Dyscalculia is far less investigated compared to dyslexia or autism spectrum disorder (Bishop, 2010), yet it is as prevalent as dyslexia (Butterworth et al., 2011) and autism spectrum disorder (Elsabbagh et al., 2012), affecting about 5 to 6% of the population (Rubinsten and Henik, 2009). It is a life-long disorder with serious consequences throughout life for income (Estrada-Mejia et al., 2016), socio-economic status (Ritchie and Bates, 2013), medical decision making (Reyna et al., 2009), and even mortgage default (Gerardi et al., 2013).

Dyscalculia is thought to originate from impaired numerical magnitude processing (De Smedt et al., 2013), but to date no study has investigated the neural quality of these magnitude representations and their access in adults with dyscalculia. The available neuroimaging studies in dyscalculia, which mostly involve studies in children, typically considered only the overall activation level in cortical regions associated with number processing and restricted their focus largely to the intraparietal sulcus (IPS). These studies have revealed mixed results and, depending upon task requirements, hypo-activation (Ashkenazi et al., 2012; Mussolin et al., 2010b; Price et al., 2007) as well as hyper-activation (Rosenberg-Lee et al., 2015; Simos et al., 2008) in the IPS has been reported. These differences in brain activity suggested an inappropriate task-modulation of the IPS during number processing, yet they did not directly provide any information about the quality of the involved representations.

On the other hand, it has also been suggested that the number representations themselves are not impaired, but that these representations are difficult to access (Noël and Rousselle, 2011) (Figure 1a). This is supported by behavioral studies showing that children with dyscalculia were only impaired in processing symbolic but not non-symbolic magnitudes (De Smedt and Gilmore, 2011; Rousselle and Noël, 2007). At the neural level, there is some preliminary evidence for impaired access and connectivity. In children with dyscalculia reduced white matter tracts in right temporal-parietal areas (Rykhlevskaia et al., 2009), and hyper-connectivity between the IPS and lateral fronto-parietal regions, and between the IPS and the default mode network (Rosenberg-Lee et al., 2015) have been reported.

**Figure 1.**
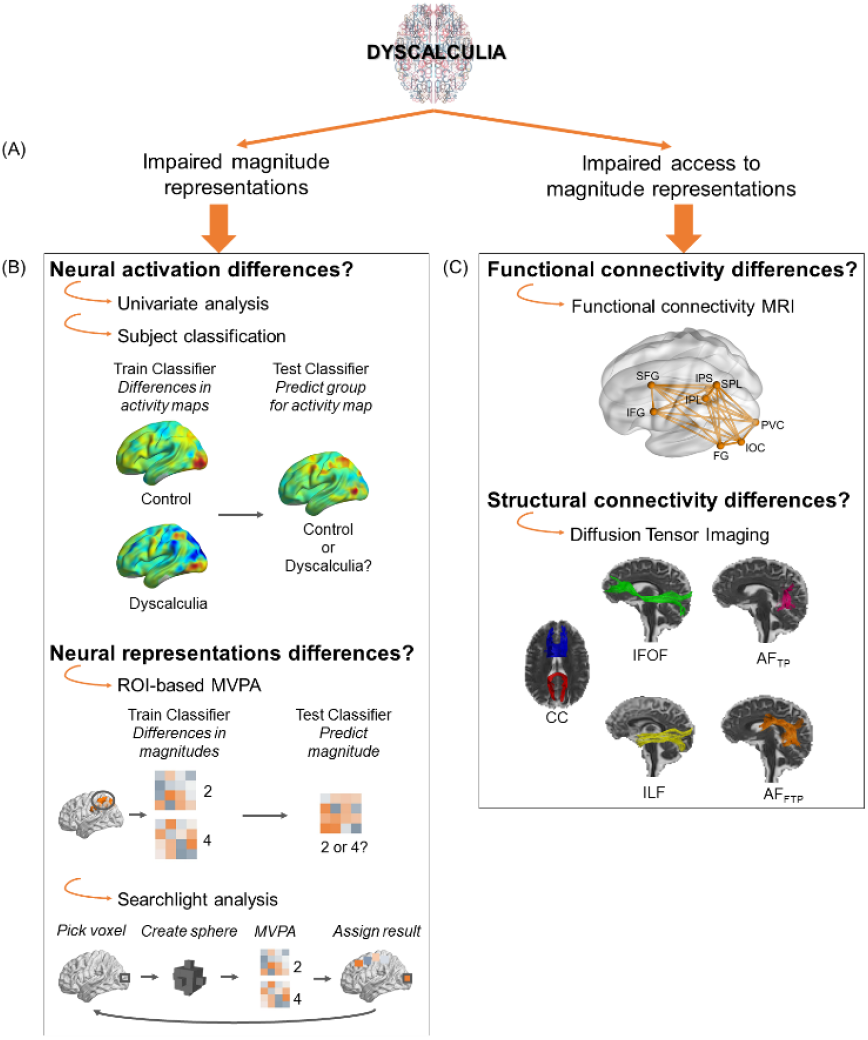
Overview of the current study. Two suggested hypotheses about the etiology of dyscalculia (*A*), and the main approaches to test them (*B-C*). Abbreviations: Regions of Interest (ROI), multivoxel pattern analysis (MVPA), magnetic resonance imaging (MRI), superior and inferior frontal gyrus (SFG, IFG), intraparietal sulcus (IPS), inferior parietal lobule (IPL, SPL), fusiform gyrus (FG), inferior occipital cortex (IOC), primary visual cortex (PVC), corpus callosum (CC), inferior fronto—occipital fasciculus (IFOF), inferior longitudinal fasciculus (ILF), tempo-parietal arcuate fasciculus (AF_TP_), and fronto-temporoparietal arcuate fasciculus (AF_FTP_).

The current study is the first to directly test both the quality of neural magnitude representations in dyscalculia as well as the access to these neural representations. We therefore coupled multivoxel pattern analysis (MVPA) (Figure 1b) with measures of functional and structural connectivity (Figure 1c). Starting from the study that tested similar hypotheses on the causes of dyslexia (Boets et al., 2013), we extended the spectrum of analyses even further by additionally including subject classification methods as well as voxel-based morphometry analyses. Subject classification methods allowed us to directly test which functional differences between the two groups were strong enough to identify individuals with dyscalculia based only on their brain activity or structure. Voxel-based morphometry analysis was considered in view of the anatomical abnormalities, for example in IPS, that have been reported in children with dyscalculia (Rotzer et al., 2008; Rykhlevskaia et al., 2009).

## 2. Material & Methods

### 2.1. Participants

In total, 54 adult participants took part in this study as paid volunteers. Due to technical issues with the scanner, a useful dataset was only acquired for 48 participants (all females, aged between 18 and 27, three left-handed participants with dyscalculia, and two left-handed participants in the control group), including 24 participants with dyscalculia and 24 control participants with normal achievement in mathematics. All participants had normal or corrected-to-normal vision and reported no neurological or psychiatric history. An interview with all the participants was conducted to confirm that all individuals with dyscalculia and none of the control participants met the DSM-V criteria for dyscalculia. All participants provided two written informed consents, one before the behavioral session and one prior to scanning. The study was approved by the medical ethics committee of KU Leuven.

### 2.2. Matching dyscalculia group and control group

All participants successfully completed the secondary school level and were either in college or university. The two groups were individually matched pairwise for their education in secondary school and college/university, gender, and age (Table 1). We evaluated their arithmetic and reading skills, motor speed, and intelligence to ensure a matching between the two groups for all measures, besides arithmetic. Statistical analyses for all the behavioral measures were done in Matlab version 8.3.0.532 (R2014a).

**Table 1.** Descriptive statistics on matching variables.

|  | Dyscalculia group | Control group | *t*[46] | *p* |
| --- | --- | --- | --- | --- |
| **Descriptive information** |  |  |  |  |
| N | 24 | 24 | - | - |
| Age (in years) | 21.96 (2.16) | 21.67 (2.20) | 0.46 | 0.65 |
| **Mathematical abilities** |  |  |  |  |
| French Kit | 35.00 (7.97) | 54.50 (15.86) | 5.38 | <0.0001 |
| Tempo Test Arithmetic | 113 (17.46) | 148.75 (21.92) | 6.25 | <0.0001 |
| Arithmetic (WAIS)<sup>a</sup> | 7.50 (2.06) | 10.79 (2.21) | 5.34 | <0.0001 |
| **Reading** |  |  |  |  |
| Z-score Reading | -0.20 (1.12) | 0.20 (0.62) | 1.49 | 0.14 |
| **IQ measures** |  |  |  |  |
| Nonverbal– Matrix reasoning (WAIS) <sup>a</sup> | 9.08 (2.90) | 10.23 (3.01) | 1.32 | 0.19 |
| Verbal – Vocabulary (WAIS) <sup>a</sup> | 10.79 (2.99) | 12.08 (2.26) | 1.69 | 0.10 |
| **Motor speed task** |  |  |  |  |
| Accuracy (% correct) | 96.88 (10.82) | 98.75 (2.21) | 0.83 | 0.41 |
| Reaction time (s) | 0.40 (0.08) | 0.36 (0.06) | -1.80 | 0.08 |

Note: Standard deviations are shown in parentheses. <sup>a</sup> Standardized score with *M* = 10 and *SD* = 3.

First, differences in mathematical abilities were assessed by three tests. Tempo Test Calculation (De Vos, 1992) and French Kit (French et al., 1963) are two standardized paper-and-pencil tests for one- and multi-digit calculation (addition, subtraction, multiplication, and division) under time pressure. Additionally, the WAIS (Wechsler Adult Intelligence Scale III) arithmetic subtest, which involves the solution of verbally presented word problems without time pressure, was administered for each participant.

Second, reading abilities were assessed by reading as many existing words (Brus, 1999) and pseudo-words (Van den Bos, 1999) as possible in one minute. This was additionally done to further verify the absence of comorbidity with dyslexia.

Third, measures of verbal and spatial intelligence were obtained by means of the, respectively; Vocabulary and Matrix Reasoning subtests of the WAIS.

Lastly, to ensure that any group differences in reaction time for mathematics or number processing tasks in the scanner were not due to group differences in processing speed, participants performed a motor speed task on a computer. During this task subjects had to decide, in a quick but accurate manner, on which side the stimulus with a white surface was presented, by pressing the corresponding key (De Smedt and Boets, 2010).

Table 1 presents a summary of the results of all these tasks, and all participants’ scores were in a normal range for the non-numerical tasks. Two-sample *t*-tests showed that groups did not significantly differ on any of the measured factors, except for the tests measuring mathematical abilities. This all confirms an appropriate matching between the two groups.

### 2.3. fMRI data acquisition and analyses

#### A. Stimuli

The stimuli and design in this experiment were the same as a previous study (Bulthé, De Smedt, & Op de Beeck, 2014). Stimuli in the experimental runs consisted of the numerical magnitudes 2, 4, 6 or 8, either displayed as symbolic numbers or as a collection of white dots on a black background (non-symbolic numbers). We controlled the stimuli for intensive (individual item size and inter-item spacing) and extensive (total luminance and total area spanned by the non-symbolic numbers) confounding parameters by varying them randomly across the dot displays (Dehaene et al., 2005). Adaptation of the symbolic numbers was minimized by varying the position and size across trials.

Stimuli were presented via Psychtoolbox 3 (Brainard, 1997) and via a NEC projector projected onto a screen located approximately 46 cm from participants’ eyes.

#### B. Design

The experimental runs had a short-block design with variable block duration (same design as in Bulthé et al., 2014). Short blocks (4, 5 or 6s) were used to prevent loss of attention and to minimize potential adaptation effects during one condition. A fixation block of 8s was presented at the beginning and end of the run. During the duration of one experimental block the same numerosity in the same format was repeated in sequences of 4, 5 or 6 trials. During the experimental runs, participants had to perform a number comparison task (indicate smaller or larger than five) every time the numerosity and/or format changed, which made the participants explicitly access numerical magnitude representations (Piazza et al., 2004). Per participant, between 8 and 12 experimental runs were acquired.

Statistical analyses for the behavioral measures (accuracy and reaction time) of the number comparison task in the scanner were done in Matlab version 8.3.0.532 (R2014a).

The localizer runs consisted of the same design as (Bulthé et al., 2015, 2014a). Participants had to subtract two numbers ranging from 1 to 20 from each other and needed to indicate if the solution was even or odd. Two localizer runs were obtained per participant.

#### C. fMRI Data Acquisition

fMRI data was acquired in a 3T Philips Ingenia CX Scanner with a 32-channel head coil using a T2* -weighted echo-planar (EPI) pulse sequence (50 slices, 2.10 × 2.15 mm in plane acquisition voxel size, slice thickness 2 mm, interslice gap 0.2 mm, repetition time (TR) = 3000 ms, echo time (TE) = 30 ms, flip angle = 90, 100 × 97 acquisition matrix). For each participant also a T1-weighted anatomical volume was obtained (182 slices, resolution 0.98 × 0.98 × 1.2 mm, TR = 9.6 ms, TE = 4.6 ms, 256 × 256 acquisition matrix).

#### D. fMRI Preprocessing

The data were processed using the Statistical Parametric Mapping software (SPM 12, Wellcome Department of Cognitive Neurology, London) in Matlab. Anatomical images were normalized to the standard brain template defined by the Montreal Neurological 152-brains average. Functional images were corrected for slice timing differences. Realignment between images to correct for motion across and within sessions was done, resulting in a set of motion parameters that were used as confounds when modelling the general linear model. No runs for any of the participants had to be excluded for extensive motion (based on a criterion of movement in any direction for more than one voxel size). Coregistration of the functional data and the anatomical image was performed. During normalization functional images were resampled to a voxel size of 2×2×2 mm. Functional images were spatially smoothed to suppress high-frequency noise by convolving them with a Gaussian kernel of 4 mm full-width at half maximum (FWHM) for subsequent multivariate voxel pattern analyses and 8 mm FWHM for subsequent second-level univariate analyses.

#### E. Statistical Analysis

For each voxel the experimental effect in a block was estimated by applying a general linear mode. This resulted in beta-values for each condition (including the fixation condition) and six motion parameters per run. *T*-statistics (resulting from conditions vs. baseline) were estimated and used as input for subsequent multivariate analysis as *t-*statistics take both the mean and the variance of the activations into account (Misaki et al., 2010). For the IPS, the analyses were repeated using the beta values, resulting in very similar effects.

#### F.ROIs

We selected ROIs that were activated during number comparison tasks in previous studies (Bulthé et al., 2015, 2014a; Holloway et al., 2013; Lyons and Ansari, 2009; Piazza et al., 2007). We selected ROIs on 4 “spatial scale” levels (Figure 2): (1) All regions (all grey matter voxels with significant activity task vs. baseline in the localizer task in a participant), (2) The 4 lobes: frontal cortex, parietal cortex, temporal cortex, occipital cortex, (3) Specific regions: fusiform gyrus (FG), inferior frontal gyrus (IFG), superior frontal gyrus (SFG), inferior occipital gyrus (IOG), superior occipital gyrus, primary visual cortex (PVC), supramarginal gyrus, angular gyrus, inferior parietal lobule (IPL), superior parietal lobule (SPL), IPS, (4) Sub-regions within the IPS: left anterior IPS (IPS_LA_), right anterior IPS (IPS_RA_), left posterior IPS (IPS_LP_), and right posterior IPS (IPS_RP_).

**Figure 2.**
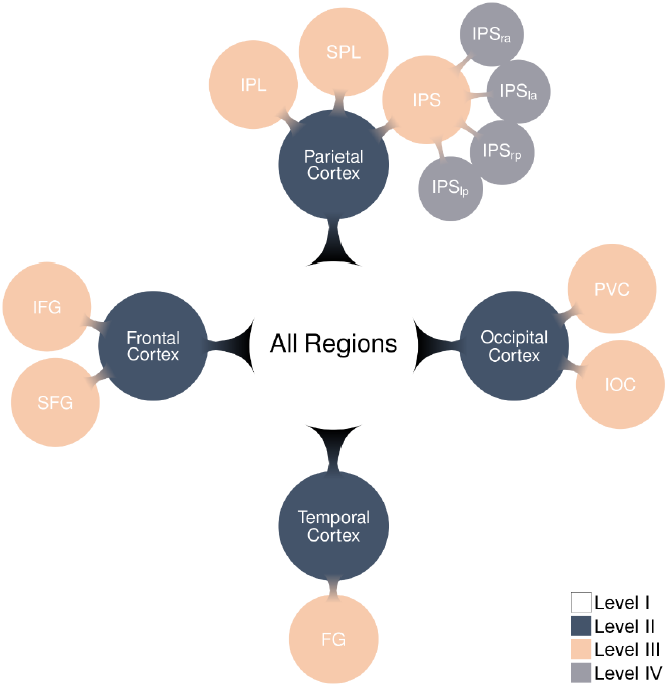
Overview of the hierarchical structure for the ROIs. ROIs that belonged in the same level are indicated by the same color. Only in the case of a significant group effect in the higher level, the lower layer was tested for both groups. Each time, FDR correction was applied for all the ROIs belonging to the same level. Abbreviations: left anterior IPS (IPS_la_), right anterior IPS (IPS_ra_), left posterior IPS (IPS_lp_), and right posterior IPS (IPS_rp_).

For those ROIs for which an anatomical mask was available in the WFU PickAtlas Toolbox (Wake Forrest University PickAtlas, fmri.wfubmc.edu/cms/software), we selected the voxels based on the conjunction of the voxels in that mask surviving the functional contrast (task minus fixation) from the independent localizer scans at an uncorrected threshold at p < 0.001. For the IPS and its subparts there was no anatomical mask available, so we delineated these ROIs manually on the functional contrast of the localizer scans (uncorrected threshold at *p* < 0.01). All ROIs were created at individual level.

The following ROIs were excluded from further analyses because they could not be defined in at least five subjects: supramarginal gyrus, superior occipital gyrus, and angular gyrus. For other ROIs, if it was not possible to define the ROI in a participant, this participant was excluded in subsequent analyses for that ROI. There were no discrepancies between how many ROIs could or could not be defined in participants between the two groups.

#### G. Correction for multiple comparisons

We applied the same hierarchical FDR-correction as in a previous study (Bulthé et al., 2014a). All group differences, for univariate and multivariate ROI-based analyses, were tested with a two-sided two-sample *t-*test. In a first step, the group difference was tested for both formats in the All Regions ROI (level 1). If there was a significant difference between the two groups for a format in the All Regions, we tested whether there was a significant group difference in the four lobes (level 2) for that format. For the lobes showing a significant group difference, the smaller ROIs (level 3) in that lobe were subsequently tested with the following sublevels per lobe: (temporal) fusiform gyrus, (frontal) inferior frontal gyrus, superior frontal gyrus, (occipital) inferior occipital gyrus, superior occipital gyrus, primary visual cortex, (parietal) supramarginal gyrus, angular gyrus, inferior parietal lobule, superior parietal lobule, and IPS. If the IPS showed a significant group difference, the subparts of the IPS (level 4) were tested. Within each level of the hierarchy a correction for multiple comparisons (false discovery rate, FDR) across all ROIs in that level was applied. This reasoning is similar to the well-known statistical approach of only testing a priori *t*-contrasts (e.g., pairwise comparisons) if an *F*-test including all conditions is significant. To avoid to have missed any important smaller ROIs because a higher level ROI did not show a significant group difference, we also applied a FDR-correction across all ROIs. However, never did any of the ROIs survived this more stringent FDR-correction if the higher-level ROI did not show an effect.

#### H. Univariate analysis

For every participant two contrasts for the experimental runs were estimated: symbolic numbers minus fixation and non-symbolic numbers minus fixation. For this analysis, no distinction was made between the different numerical magnitudes within each format.

A second-level group analysis in SPM12 was done for these contrasts to test for activation differences for symbolic numbers and non-symbolic numbers between the two groups on a whole brain level (threshold of *p* < 0.05 after FWE correction at voxel-wise level). Figures for this analysis were made with BrainNet Viewer (Xia et al., 2013).

A ROI-based univariate analysis was also conducted to test for group differences within each format. We performed two-sample *t*-tests and corrected for multiple comparisons in a hierarchical manner, as described above.

#### I. Multivariate analysis

##### Subject classification based upon spatial variation in univariate activity levels

Instead of decoding different conditions within one subject, it is possible to investigate whether we can decode between the functional data of participants from the control group and the dyscalculia group. In other words, can these two groups be differentiated on the basis of their functional brain activity? For this analysis, we used the functional contrasts ‘symbolic numbers minus fixation’ and ‘non-symbolic numbers minus fixation’ from the experimental runs for every participant. It is important to point out that this analysis does not tackle the underlying differences in the quality of the neural representations of symbolic numbers and non-symbolic numbers between the groups, but tests if there is a more general difference in activation between the two groups when symbolic and non-symbolic numbers are processed. In this way, this analysis is more closely related to a second level univariate analysis than to multivariate ROI-based decoding and searchlight analysis. The main difference with a second level univariate analysis is that there is no activity-based comparison at the level of single voxels, but a spatial pattern comparison between the two groups across the whole brain or across a selected ROI.

The classification was performed with linear support vector machines with the following parameters: a radial basis function kernel as decision function with parameter gamma set to 1; a C-SVC classification algorithm was used with parameter C set to 1. We applied a leave-one-pair-out-cross-validation (LPOCV) technique, similar to the one used in Ung *et al.* (2014). With this method the classifier was trained on all participants, except for one random participant from the control group and one random participant from the dyscalculia group. Afterwards, the trained model was tested on this left-out pair. This procedure was repeated until each participant was left out once. Because of this random division into pairs, slightly different accuracies can occur depending on the division. For this, the LPOCV was run 1000 times, and the results were averaged across all these repeats.

Statistics were obtained by a Monte Carlo Permutation test (Mourã o-Miranda et al., 2005). The class labels of the training set were 1000 times randomly permuted and the same LPOCV procedure as described above was applied. A p-value for the subject classification accuracy was obtained by the number of times the permutation accuracy is greater than or equal to the subject classification accuracy, divided by 1000. For this subject classification, we noted that the permutation-based threshold for statistical significance was actually very similar to the threshold as it would been set by a simple parametric binomial test taking into account the proportion of participants classified in a particular group. To correct for multiple comparisons, we applied a FDR-correction across the four lobes and across the IPS regions.

##### Whole brain searchlight analysis of the decoding of neural magnitude representations

The method is particularly suited for finding where in the brain the local spatial activity pattern differs across conditions without selecting any ROIs (Kriegeskorte et al., 2006; Kriegeskorte and Bandettini, 2007).

For the searchlight analysis, we used “The Decoding Toolbox” together with own custom made code in Matlab (Hebart et al., 2015). The classification model (SVM) and its parameters for the ROI-based decoding were similar to the ones used for the subject classification analysis.During the searchlight analysis a sphere with a radius of two voxels (volume of max. 33 voxels) was sequentially moved across the entire grey-matter volume (similar as Bulthé *et al.* (2014)). The searchlight analysis resulted in a map for each format per participant. Afterwards, the maps were spatially smoothed using Gaussian kernels of 8 mm FWHM (equal to the univariate smoothing level). Finally, a second-level analysis was done in SPM12 to test for group differences for both formats (threshold of *p* < 0.05 after FWE correction).

##### ROI-based decoding of neural magnitude representations

For each ROI, a decoding classification analysis was implemented with custom code written in Matlab (The MathWorks, Natrick, MA) using the LIBSVM toolbox (Chang and Lin, 2011). The classification model (SVM) and its parameters for the ROI-based decoding were similar to the ones used for the subject classification and searchlight analyses.

Response patterns for every condition in each run were extracted for each ROI, with in each pattern the *t*-values of all voxels in the ROI. The patterns were standardized by subtracting the mean across voxels and then dividing this by the standard deviation across voxels for each condition. We followed a repeated random subsampling cross-validation procedure: The data were randomly divided into 70% training data and 30% test data (the latter were averaged to one response pattern per condition), and this was repeated 100 times.

The decoding accuracies were then averaged over two comparisons of interest: symbolic numbers (mean within-format decoding accuracy for symbolic numbers) and non-symbolic numbers (mean within-format decoding accuracy for non-symbolic numbers).

Group differences were tested with a two-sample *t*-test and corrected for multiple comparisons with FDR for the four lobes and for the IPS subparts separately.

#### J. Functional connectivity analysis

Preprocessing steps for this analysis comprised (1) bandpass filtering between 0.01 and 0.2 Hz (Balsters et al., 2016; Baria et al., 2013), (2) regression of head motion parameters and their first derivatives (3) regression of white matter and ventricle signals and their first derivatives (Ebisch et al., 2013), (4) regression of task-related BOLD fluctuations (task = the contrast ‘task minus baseline’) (Boets et al., 2013; Ebisch et al., 2013), (5) scrubbing of motion-affected functional volumes (Power et al., 2012), and (6) spatial smoothing at 4 mm FWHM. The eight ROIs of level III, which were created for each participant for above analyses, were included as seed ROIs for the functional connectivity analysis. We obtained a representative BOLD time course for each ROI by averaging the time courses of the voxels within the ROI. For each participant we then created a functional connectivity matrix by calculating Pearson cross-correlations between the BOLD time courses of each pair of ROIs. After converting the single-subject matrices to Z-scores by means of the Fisher’s r-to-Z transformation, we calculated a group-level matrix by conducting a random effects analysis across subjects (*p*_FDR_ < .001). Group-level comparisons among functional connectivity scores were performed by calculating independent-sample *t* tests on the Z-score matrices (*p*_FDR_ < .05).

### 2.4. DTI data acquisition and analyses

#### A. DTI data acquisition

For 23 control participants and 21 participants with dyscalculia DTI data were obtained. Diffusion images were acquired on a 3T Philips Ingenia CX Scanner using a single spin shot EPI with SENSE acquisition. Whole brain images were acquired with the following parameters: 58 sagittal slices, slice thickness = 2.5 mm, voxel size = 2.5×2.5×2.5 mm^3^, repetition time = 7600 ms, echo time = 82 ms, field-of-view = 220 × 240 mm^2^, matrix size = 80 × 94 and acquisition time = 10 min 32 s. Diffusion gradients were applied along 60 noncollinear directions (*b =* 1500 s/mm^2^).

#### B. Image preprocessing and tractography

Preprocessing of the raw diffusion MR data was done using ExploreDTI (Leemans et al., 2009) and contained following steps: (1) Images were corrected for eddy current distortion and subject motion; (2) a non-linear least square method was applied for diffusion tensor estimation, and (3) for each participant a whole brain tractography was estimated using following parameters: uniform 2 mm seed point resolution, FA threshold of 0.2, angle threshold of 40°, and fiber length range of 50 – 500 mm.

We used the TrackVis software to delineate white matter tracts for each participant in native space (Wang and Wedeen, 2007). Against the background of the review of Matejko and Ansari (2014), the following tracts were delineated: genu and splenium of the corpus callosum (CC), left and right inferior fronto-occipital fasciculus (IFOF), left and right inferior longitudinal fasciculus (ILF), left and right frontal to temporoparietal arcuate fasciculus (AF_FTP_), left and right frontal to temporal AF (AF_FT_), left and right frontal to parietal AF (AF_FP_), and left and right temporal to parietal AF (AF_TP_). For each of the tracts the fractional anisotropy (FA), mean diffusivity (MD), axial diffusivity (AD), and radial diffusivity (RD) values were extracted for every participant. The delineation of all the tracts for each participant of the control group was done by two independent raters. The inter-rater reliability for all tracts ranged from 0.87 to 0.99, demonstrating a high reproducibility of the tractography.

To test for the difference in white matter connectivity between the two groups, two sample *t-* tests were performed in Matlab for each DTI measures (FA, MD, AD, and RD values) of each tract. For each DTI measure, a FDR correction for multiple comparisons was applied across tracts.

### 2.5. Voxel Based Morphometry (VBM)

The VBM analysis was performed with SPM12 and according to the methodological description of Ashburner & Friston (2000) for the standard VBM analysis (Good et al., 2001). The structural MRI images of all participants were spatially normalized to Talairach space and resliced to a voxel size of 1 mm^3^ isotropic. The resliced images were partitioned into grey matter, white matter, cerebrospinal fluid, and other compartments. Grey matter segments were smoothed with a 12-mm FWHM isotropic Gaussian kernel. Statistical analysis for comparing grey matter volume between the two groups was performed by a two-sample *t*-test with global normalization for total amount of grey and white matter (threshold of *p* < 0.05 after FWE correction).

## 3. Results

Behavioral and neuroimaging data were collected in 24 college/university students with and 24 college/university students without dyscalculia. All adults with dyscalculia met the DSM-V (American Psychiatric Association, 2013) criteria for specific learning disorder. Both groups were matched on sex, intelligence, age, educational history and reading ability. They differed significantly, as expected, in their mathematical abilities (Table 1).

### Behavioral Analysis

A 2 × 2 ANOVA (group x format) was performed to test for group and format differences on the number comparison task during the experimental runs. For accuracy there was no significant group difference between the control group (95.75%) and dyscalculia group (94.77%) (*F*_1,92_ = 2.29, p = 0.13). A significant effect for format was observed and accuracy was higher for symbolic numbers than for non-symbolic numbers (*F*_1,92_ = 41.04, p < 0.001): 97.35% and 93.17%, respectively. There was no significant interaction effect between group and format (*F*_1,92_ = 0.01, p = 0.92).

For reaction time, a significant group difference was observed with faster response times for controls (0.90s) than for individuals with dyscalculia (1.18s) (*F*_1,92_ = 41.2, p < 0.001). Again, a significant format effect was present with faster reaction times for symbolic numbers (0.91s) than for non-symbolic numbers (1.17s) (*F*_1,92_ = 34.67, p < 0.001). There was no interaction between group and format (*F*_1,92_ = 0.36, p = 0.55).

Thus, overall, the two subject groups performed the task equally well, but individuals with dyscalculia were significantly slower compared to the control group. This allowed us to directly compare our findings with the earlier study on dyslexia as a similar behavioral pattern was found (Boets et al., 2013).

### 3.1. Neural activation levels for symbolic and non-symbolic numbers

#### A. Univariate analyses

In a first step, in line with earlier studies, we performed univariate analyses to test for group differences in overall activation level. No significant group differences for symbolic and non-symbolic numbers (Figure S1) were found on a whole brain voxel-wise *t*-test (second-level analysis, voxel-wise FWE corrected at *p* < 0.05). This is, on average, consistent with earlier observed task-dependent hyper- and hypo-activations in dyscalculia.

We additionally performed a ROI-based univariate analysis with two categories of ROIs. We selected ROIs in the parietal and frontal cortex which have been frequently related to numerical processing (Ansari, 2008; Feigenson et al., 2004). On the other hand, we included ROIs in the occipital and temporal cortex that are often studied in other relevant research domains (visual perception and reading) but that are not typically correlated with numerical processing.

The 17 ROIs were tested in a hierarchical order as introduced before in Bulthé et al. (2014a) (Figure 2). This approach combines the strength of global large-scale volumes to pick up highly distributed effects with the strength of smaller local ROIs to pick up more focal differences (Bulthé et al., 2014b). Four spatial scales were introduced: whole cerebral cortex (“All Regions ROI(”), lobes, regions, and sub-regions. At the first level, we checked for group differences across all grey matter. If (and only if) there was a significant group difference at a higher level, the group comparison was tested for the lower level and *p*-values were FDR corrected across that level.

For the ROI-based univariate analysis, we found no significant differences in brain activity between controls and adults with dyscalculia for either formats in the All Regions (Dots: *t*_46_ = 1.44, *p* = 0.11; Arabic digits: *t*_46_ = 1.86, *p* = 0.07). There were also no significant differences in activity in any of the individual ROIs (lowest *p*_*FDR*_ = 0.10).

### B. Subject classification

To further investigate whether there were distinguishable patterns of activation versus fixation between the two groups, we applied a subject classification procedure (see Methods). Subject classification allowed us to examine if the activation patterns of both groups for either formats at various spatial levels were different enough to be picked up by a classifier.

The results of the subject classification analysis did not show a significant subject classification accuracy using the patterns in the All Regions ROI, for non-symbolic (classification accuracy = 0.57, *p* = 0.17) nor for symbolic (classification accuracy = 0.51, *p* = 0.45) numbers. None of the ROIs showed a significant effect either at an FDR-corrected level for any of the formats. Thus, even with sensitive classification methods we did not find significant differences between subject groups in terms of the general pattern of activation. These analyses suggest that the same representations and processes seemed to be involved in the two groups.

### 3.2. Quality of neural representations of symbolic and non-symbolic numbers

#### A. Searchlight analysis

The above described results showed no significant group differences in level of activation for either formats. All these analyses are based upon the level of activation versus fixation. Earlier studies also focused upon such activation levels. Here we proceed with more refined analyses which allow to assess the quality of neural representations. Therefore, we applied a whole-brain MVPA searchlight analysis based upon the decoding of different magnitudes. This analysis was performed separately for the two formats, non-symbolic and symbolic magnitude representations.

The searchlight analysis demonstrated specific (‘hotspots(’ in both groups for non-symbolic and symbolic numbers representations (Figure 1, first two rows). For control participants the non-symbolic representations of different numerosities (e.g., 4 dots versus 8 dots) were distinguishable in many regions of the dorsal stream. In individuals with dyscalculia, the non-symbolic number representations were distinct in the occipital pole and a few patches in the parietal cortex (mainly superior parietal lobule).

An explicit comparison between groups for non-symbolic numbers (Figure 3) clearly demonstrated significantly less distinct non-symbolic number representations in the anterior parietal, frontal lobe, and a small spot in the temporal lobe for individuals with dyscalculia compared to controls.

**Figure 3.**
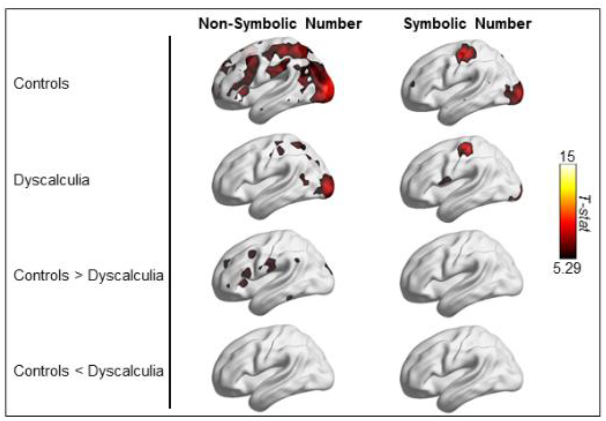
Searchlight results. Illustration of the decoding accuracies elicited by non-symbolic and symbolic numbers in the both groups. The difference in decoding accuracies between the two groups for each format is also illustrated. The results were corrected with FWE (*p* < 0.05).

For symbolic numbers we found much less regions with distinct number representations for both groups compared to non-symbolic numbers. This is consistent with earlier studies using a similar paradigm and data-analytic methods (Eger *et al.*, 2009; Damarla & Just, 2012; Bulthé, *et al.*, 2014) which also showed a much lower ability to decode symbolic numbers compared to non-symbolic numbers. The searchlight maps of both groups were not significantly different for symbolic numbers, and thus it seems that for symbolic numbers there is no difference in how overlapping the neural representations are between both groups.

In both groups the classifier was also able to distinguish between symbolic numbers in the occipital pole and motor and somatosensory cortex. The significant accuracy in the motor and somatosensory cortex might be due to a response confound. Most (four out of six) pairwise comparisons of numbers are between conditions triggering different motor responses (e.g. smaller or larger than 5) and this might have been picked up by the classifier. Note that in a different study with the same paradigm and only sixteen subjects we did not observe this effect (Bulthé et al., 2014a), suggesting a small effect that can only be picked up by the classifier with enough data. These regions did not show any group difference.

#### B. ROI-based decoding

Importantly, the searchlight results should not be interpreted as evidence that number representations are relatively focal, because searchlight analyses are notoriously biased towards finding focal representations (Bulthé et al., 2014b). Therefore, we also applied ROI-based decoding analyses to look for more widespread differences between the two groups, first on larger spatial scale (All Regions and different lobes), subsequently moving towards smaller spatial scales (IPS, temporal, frontal, and occipital regions).

To test if there were distinct underlying neural representations for symbolic and non-symbolic numbers in each ROI, a classifier was trained and tested to differentiate between the different numerical magnitudes within one format for a certain ROI. If there were distinct neural representations in that ROI, the decoding accuracies should be significant, with higher decoding accuracies indicating more distinct neural representations. We also compared the decoding accuracies of both groups for each format.

##### All Regions

The classifier was able to distinguish the neural representations of non-symbolic (controls: *t*_23_ = 21.59, p < 0.001; dyscalculia: *t*23 = 17.37, p < 0.001) and symbolic numbers (controls: *t*_23_ = 4.50, p < 0.001; dyscalculia: *t*_23_ = 5.89, p < 0.001) in both groups (Figure 4). For non-symbolic numbers, the neural representations were more separable in controls than in dyscalculia (*t*_46_ = 2.16, *p* = 0.04). For symbolic numbers there was no significant difference in the quality of the neural representations between controls and dyscalculia (*t*_46_ = −0.56, *p* = 0.58). Given that decoding performance was markedly lower for symbolic than for non-symbolic numbers, with decoding performance going down from 63% to 47% overall, this lack of a significant group difference for symbolic numbers might arise from a lack of sensitivity to detect a possible underlying group difference.

**Figure 4.**
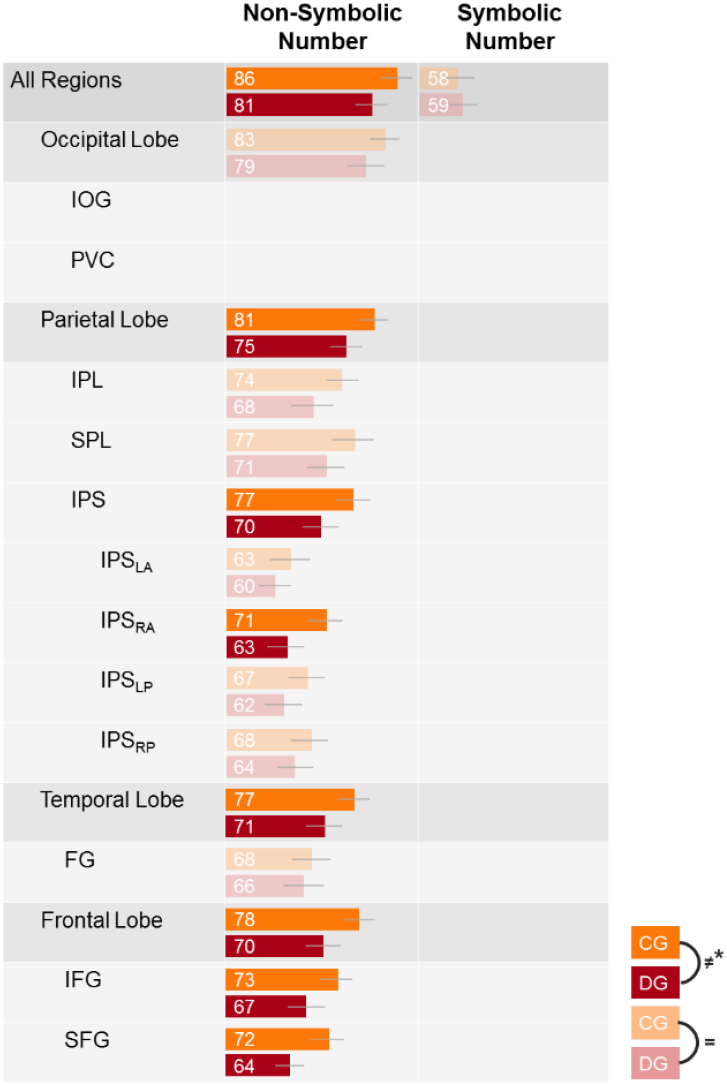
Decoding accuracies of controls and dyscalculia for non-symbolic and symbolic numbers for every ROI in hierarchical order. Orange represents the control group (CG) and red the group with dyscalculia (DG). All the within-group decoding accuracies were significantly different from chance level (FDR corrected). The dark colored bars represent a significant group difference between controls (orange) and dyscalculia (red) (FDR corrected). If a higher order ROI did not show a significant group difference (dimmed bars), lower ROIs were not analyzed and not shown in the figure. Error bars represent the 95% confidence interval for the decoding accuracy for that group in that ROI.

##### Lobes

The decoding accuracies for non-symbolic numbers were significantly different from chance level for both groups in all lobes: occipital (controls: *t*_23_ = 21.22, *p*_*FDR*_ < 0.001; dyscalculia: *t*_23_ = 14.52, *p*_*FDR*_ < 0.001), parietal (controls: *t*_23_ = 19.66, *p*_*FDR*_ < 0.001; dyscalculia: *t*_23_ = 14.61, *p*_*FDR*_ < 0.001), temporal (controls: *t*_23_ = 15.55, *p*_*FDR*_ < 0.001; dyscalculia: *t*_23_ = 10.75, *p*_*FDR*_ < 0.001), and frontal (controls: *t*_23_ = 16.10, *p*_*FDR*_ < 0.001; dyscalculia: *t*_23_ = 10.94, *p*_*FDR*_ < 0.001). There was no significant group difference in the occipital lobe (*t*_46_ = 1.63, *p*_*FDR*_ = 0.11). A significant group difference was observed in the parietal lobe (*t*_46_ = 2.49, *p*_*FDR*_ = 0.03), distinguishable neural patterns for non-symbolic numbers in controls than in dyscalculia.

##### Parietal ROIs

The neural patterns for non-symbolic numbers were distinct from each other in each group in the IPL (controls: *t*_23_ = 14.05, *p*_*FDR*_ < 0.001; dyscalculia: *t*_23_ = 8.07, *p*_*FDR*_ < 0.001), SPL (controls: *t*_23_ = 12.22, *p*_*FDR*_ < 0.001; dyscalculia: *t*_23_ = 10.45, *p_FDR_* < 0.001), and the IPS (controls: *t*_23_ = 13.91, *p*_*FDR*_ < 0.001; dyscalculia: *t*_23_ = 10.05, *p*_*FDR*_ < 0.001). Although there was a strong trend for more distinguishable neural patterns in controls than dyscalculia in the IPL (*t*_46_ = 2.10, *p*_*FDR*_ = 0.052) and SPL (*t*_46_ = 1.99, *p*_*FDR*_ = 0.052), the group difference, with more distinct neural patterns in controls than dyscalculia, was only significant in the IPS (*t*_46_ = 2.50, *p*_*FDR*_ = 0.048).

##### *IPS* subparts

Because there was a significant group difference for non-symbolic numbers in the IPS, its subparts were also analyzed for presence of distinct non-symbolic numbers representations for each group. We also tested whether there was a group difference in the distinctiveness of these neural representations. In all the subparts of the IPS the classifier could reliably differentiate between the different neural representations of non-symbolic numbers for each group: IPSLA (controls: *t*_23_ = 6.22, *p*_*FDR*_ < 0.001; dyscalculia: *t*_23_ = 5.72, *P*_*FDR*_ < 0.001), IPSRA (controls: *t*_22_ = 10.86, *p*_*FDR*_ < 0.001; dyscalculia: *t*_22_ = 6.37, *p*_*FDR*_ < 0.001), IPS_LP_ (controls: *t*_23_ = 8.71, *p*_*FDR*_ < 0.001; dyscalculia: *t*_23_ = 6.09, *p*_*FDR*_ < 0.001), and IPS_RP_ (controls: *t*_23_ = 9.02, *p*_*FDR*_ < 0.001; dyscalculia: *t*_23_ = 7.49, *p*_*FDR*_ < 0.001). However, only in the IPS_R_ A the distinctiveness of the neural representations was significantly higher for controls than for dyscalculia (*t*_44_ = 2.91, *p*_*FDR*_ = 0.02).

##### Temporal ROIs

The FG contained distinct neural representations for non-symbolic numbers in controls (*t*_23_ = 8.84, *p*_*FDR*_ < 0.001) and dyscalculia (*t*_23_ = 7.53, *p*_*FDR*_ < 0.001). There was no difference in distinctiveness of these neural representations between the two groups (*t*_46_ = 0.60, *p*_*FDR*_ = 0.55).

##### Frontal ROIs

The classifier could distinguish different neural representations for non-symbolic numbers for both groups in the IFG (controls: *t*_23_ = 13.65, *p*_*FDR*_ < 0.001; dyscalculia: *t*_23_ = 8.48, *p*_*FDR*_ < 0.001) and the SFG (controls: *t*_23_ = 11.70, *p*_*FDR*_ < 0.001; dyscalculia: *t*_23_ = 8.82, *p*_*FDR*_ < 0.001). The neural representations in controls were also more distinct than in dyscalculia in the IFG (*t*_46_ = 2.50, *p*_*FDR*_ = 0.02) and the SFG (*t*_46_ = 3.35, *p*_*FDR*_ = 0.003).

### 3.3. Connectivity differences between the two groups?

We subsequently tested if the numerical impairments in dyscalculia were due to an access deficit and related connectivity differences. This was done by analyzing (A) functional connectivity between ROIs and (B) structural connectivity in specific white matter tracts (Figure 1c).

#### A. Functional connectivity

A functional connectivity analysis was performed to test which ROIs were functionally coupled with each other and whether this coupling differed between the two groups. Only the 8 ROIs on level III (Figure 2) were included. For both groups, all the pairwise functional connectivity strengths were significant (controls: 10.92 < *t*_23_ < 38.81, all *p*_FDR_’s < 0.001; dyscalculia: 12.80 < *t*_23_ < 37.58, all *p*_FDR_’s < 0.001) (Figure 5a-b). For the individual connections between ROIs, there were significant group differences for the connectivity between PVC and IOG (*t*_46_ = −3.17, *p*_FD_ R = 0.04) and between PVC and FG (*t*_46_ = −3.47, *p*_FDR_ = 0.03) with higher connectivity in individuals with dyscalculia than in controls (Figure 5c). Interestingly, none of the regions involved in this hyper-connectivity showed a group difference in the distinctiveness of neural representations in the previous analyses.

**Figure 5.**
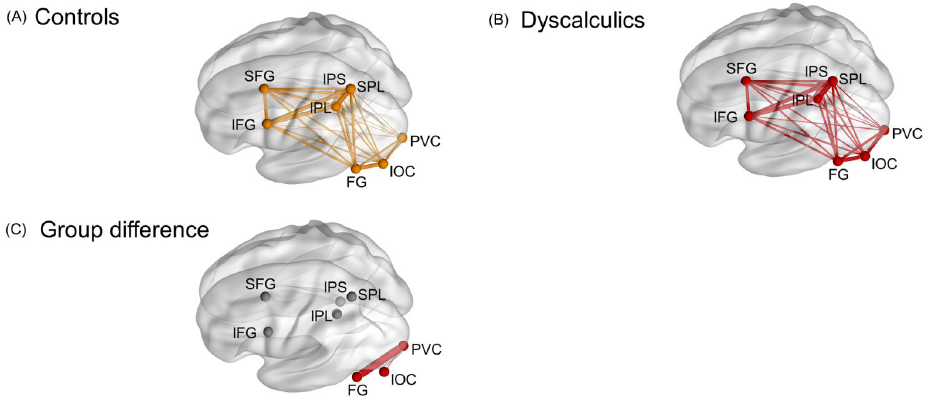
Overview of the fcMRI results. All the ROIs were significantly functionally connected with each other for the (*A*) control participants and (*B*) participants with dyscalculia at a FDR (p < 0.001) corrected level. (*C*) We observed a significant group difference at FDR (*p* < 0.05) corrected level with higher functional connectivity for dyscalculia between PVC and IOC and between PVC and FG.

#### B. Structural connectivity

DTI measures the quality of the white matter tracts that connect different regions in the human cortex. We defined the relevant tracts a priori based upon the literature. More specifically, delineated the white matter tracts that have been related to numerical and mathematical processing in previous studies, as summarized by Matejko and Ansari (2014): the genu and splenium of corpus callosum, left and right inferior fronto-occipital fasciculus, left and right inferior longitudinal fasciculus, left and right frontal to temporoparietal arcuate fasciculus (AF), left and right frontal to temporal AF, left and right frontal to parietal AF, and left and right temporal to parietal AF.

None of the tracts showed a significant group difference (even at uncorrected level for multiple comparisons) for fractional antisotropy (FA), mean diffusivity (MD), axial diffusivity (AD), and radial diffusivity (RD) values. For FA, the *t-*values ranged from −1.85 to 1.41 (0.41 < *p*_*FDR*_ < 0.91); for AD *t*-values ranged from −1.85 to 0.82 (0.54 < *p*_*FDR*_ < 0.99); for MD *t-*values ranged from −0.92 to 0.90 (all *p*_*FDR*_’s = 0.97); and for RD *t*-values ranged from −0.79 to 1.39 (0.89 < *p*_*FDR*_ < 0.98).

### 3.4. Structural differences between control and dyscalculia group?

Previous studies found decreased grey matter volume in the IPS, other parietal regions, and even frontal and temporal regions in children with dyscalculia (Rotzer et al., 2008; Rykhlevskaia et al., 2009). We performed a voxel-based morphometry analysis to investigate (i) whether the differences in the quality of neural representations between individuals with dyscalculia and controls were related to differences in grey matter volume; and/or (ii) whether any other brain regions showed structural differences.

Only an increase in grey matter in the bilateral posterior cingulate cortex (PCC) (*t*_46_ = 5.27, *p*_FWE_ = 0.04, MNI coordinates: −22 −48 25) was observed in dyscalculia compared to controls. No other differences were found. Importantly, there were no differences in grey matter volume in any of the regions that showed a difference in quality of neural representations between our two groups.

To test if and how the PCC was involved in numerical processing, we (a) verified if there were univariate activation differences between the two groups for both formats in the PCC, (b) classified subjects based on functional data (non-symbolic and symbolic numbers) within the PCC, and (c) ran a MVPA analysis for this ROI to test if distinct neural representations for non-symbolic and symbolic numbers can be detected in this area.

Univariate results demonstrated significant activation levels in PCC for symbolic numbers in controls (*t*_23_ = −2.42, *p* = 0.02), but not in dyscalculia (*t*_23_ = −1.21, *p* = 0.24) during a number comparison task. However, there were no group differences in level of activation in this area for symbolic numbers (*t*_46_ = −1.06, *p* = 0.29). Non-symbolic numbers elicited significant activation in both controls (*t*_23_ = −2.36, *p* = 0.03) and dyscalculia (t_23_ = −4.38, p < 0.001) in the PCC, but again, there was no significant group difference (*t*_46_ = −0.25, *p* = 0.80).

The subject classification for activation in the PCC by symbolic and non-symbolic numbers was not significant (classification accuracy: 0.44, *p* = 0.75; classification accuracy: 0.59, *p* = 0.10, respectively). So, the functional data related to a number comparison task in the PCC are not enough to reliably distinguish between DD and controls (in contrast to structural data in the PCC).

The MVPA analysis showed no distinct neural representations for symbolic numbers in controls (classification accuracy: 0.48, *t*_23_ = −1.48, *p* = 0.15) and dyscalculia (classification accuracy: 0.50, *t*_23_ = 0.26, *p* = 0.79). This absence of distinct neural representations for symbolic numbers in both groups makes a test for between-group differences redundant. For non-symbolic numbers, distinct neural representations could be detected in controls (classification accuracy: 0.57, *t*_23_ = 4.12, p < 0.001), but not in dyscalculia (classification accuracy: 0.53, *t*_23_ = 1.78, *p* = 0.09). However, this difference was not significant (*t*_46_ = 1.75, *p* = 0.09).

Overall, these follow-up analyses confirmed that the PCC only differed between controls and adults with dyscalculia in a *structural* way and not in functional activity or neural representations elicited by the number comparison task.

## 4. Discussion

This study combined for the first time different neuroimaging methods (anatomical, univariate and multivariate functional, and connectivity analyses) to disentangle the different hypotheses for the etiology of dyscalculia. Our results indicate impaired non-symbolic magnitude representations in the IPS, parietal regions, and other brain regions in adults with dyscalculia. This is consistent with the impaired magnitude representations hypothesis. However, we also found anatomical and functional connectivity differences in adults with dyscalculia. The latter differences are observed in brain regions which do not show a difference in the quality of magnitude representations. To conclude, the neural correlates of dyscalculia in adults cannot be reduced to only impaired magnitude representations or only impaired access to these magnitude representations, nor can they be reduced to one or a few brain areas.

The observation that the deficits in dyscalculia cannot be localized to one brain region or to one particular type of brain deficit (functional, anatomical or connectivity) has important consequences for the study of the neural basis of dyscalculia, especially because previous studies typically focused narrowly upon the IPS and parietal regions or on one specific type of analyses, often the least informative univariate analyses. Nevertheless, our findings are consistent with theoretical proposals that most neurodevelopmental disorders arise from a combination of diffuse functional disruptions, deficits in connectivity between regions, and anatomical differences (Johnson et al., 2002; Menon, 2011).

### 4.1. Functional impairments

We demonstrated that the deficits in the quality of neural magnitude representations are spread out throughout the cortex, as was already suggested based on more indirect evidence from a few studies of univariate levels of activation (Kaufmann et al., 2011; Rosenberg-Lee et al., 2015). In our study, the representations in these regions outside the parietal cortex might be related to numerical representations as well as to how these representations are recruited during more general processes. For example, frontal regions often play an important role in attention and working memory processes required for problem solving, also in numerical context.

We demonstrated this impaired quality of non-symbolic magnitude representations while participants were performing a numerical magnitude comparison task. We deliberately opted for timing parameters which allowed the participants with dyscalculia to perform the task quite well. In fact, their overall performance was not significantly different from controls, they only required more time (slower reaction time). Because overall performance was the same, the impaired quality of magnitude representations at the neural level cannot be explained by one group not being able to perform the task and hence being less motivated/attentive. If anything, the slower reaction time of the participants with dyscalculia suggests that these subjects were processing the stimuli for a longer time, which could have resulted in a better quality of the magnitude representations. Overall, the behavioral task performance avoids that we have to take into account a performance discrepancy when interpreting the neural data. Strikingly, a previous study of phonological processing in dyslexia also compared subject groups with the same accuracy and a different reaction time (slower in the dyslexic group), and in that case found no difference in the quality of neural representations (Boets et al., 2013).

While the deficit in non-symbolic number processing for dyscalculia was very clear in our study, we found no group differences in the quality of magnitude representations for symbolic numbers. This does not necessarily mean that symbolic representations are not impaired in dyscalculia. Previous MVPA studies in healthy participants already showed that these symbolic representations are harder to detect than representations of non-symbolic magnitudes (Eger *et al.*, 2009; Damarla & Just, 2012; Bulthé *et al.*, 2014; Bulthé *et al.*, 2015; Lyons *et al.*, 2015). This was confirmed in the current study as we observed smaller decoding accuracies, also in controls, for symbolic magnitudes. Given that decoding performance was markedly lower for symbolic than for non-symbolic numbers, the lack of a significant group difference for symbolic numbers might arise from a lack of sensitivity to detect a possible underlying group difference. Future studies that investigate the quality of symbolic number representations in dyscalculia might consider to collect more scanning data of only symbolic numbers. Such studies might also increase the difficulty of the task, or use larger symbolic numbers to overcome this lack of sensitivity.

Our study is the first study to demonstrate that the quality of non-symbolic magnitude representations is affected in adults with dyscalculia. Although there have been previous studies on dyscalculia, they are very different from ours because of two major differences. First, previous neuroimaging studies used univariate fMRI analyses that answer the question if there are altered levels of brain activation in individuals with dyscalculia. When we applied the same type of univariate analyses to our dataset, we failed to detect any clusters of voxels in the whole brain that showed a different level of activation in individuals with dyscalculia. It is not clear whether and in what direction our null results would deviate from earlier observed task-dependent altered levels of activations in dyscalculia as mixed results have been reported. Several studies have reported a decreased activation in dyscalculia during non-symbolic number comparison tasks (Price et al., 2007), arithmetic problem solving (Ashkenazi et al., 2012), and symbolic number comparison (Mussolin et al., 2010a). On the other hand, others have reported increased activity in the IPS during arithmetic problem solving in children with dyscalculia compared to matched controls (Rosenberg-Lee et al., 2015). Combining these previously found hyper- and hypoactivation in the IPS together, one could expect a null result, as we find.

Second, all previous functional neuroimaging work on dyscalculia has been carried out with children. Our study is the first functional neuroimaging study with adults diagnosed with dyscalculia. Hence, it was not known to date whether this functional altered activation levels in the IPS and other brain regions correlated with dyscalculia remain throughout the lifespan of persons with dyscalculia. Our results suggest that such differences in overall activity may have diminished during adulthood. This does not exclude the possibility that during development there might have been time points in development during which hyper- or hypoactivation in the IPS in children with dyscalculia is observed. On the other hand, it might be that such differences in overall activity only emerge when more complex numerical tasks (e.g. calculation instead of comparison) are used.

To conclude, our multivariate analyses results provided the first evidence that non-symbolic number representations were less precise in adults with dyscalculia in comparison to a strictly matched control group. These less precise non-symbolic number representations in dyscalculia were observed throughout the dorsal stream and the IPS, and were thus very widespread in the cortex. However, for symbolic magnitudes we did not observe any differences in neural quality between the two groups and further investigation is needed to unravel if symbolic neural representations are less precise in dyscalculia or not.

### 4.2. Connectivity deficits

The current study is the first neuroimaging study to look into the structural and functional connectivity correlates in adults with dyscalculia. We found no group differences in any of the white matter tracts (four in total with 17 segments) under investigation. However, we did observe increased functional connectivity in temporo-occipital regions.

In contrast to the functional connectivity differences in this study, we did not observe any structural connectivity differences in any of the tracts previously related to numerical processing (Matejko and Ansari, 2014). This is in contrast to two previous studies who found that children with dyscalculia had decreased connectivity in the right temporal-parietal areas (Rykhlevskaia et al., 2009), and the bilateral superior longitudinal fasciculus (Kucian et al., 2013). There are two possible explanations for the discrepancy in findings between our study and previous ones. First, we have a very strict matching between adults with and without dyscalculia, namely by controlling not only for sex, age, and intelligence, but also for educational history and environment. This strict matching can explain why we did not replicate previous neuroimaging studies. However, this strict matching is essential to dedicate the observed differences to dyscalculia and not to possible differences in education. Second, it might be that the with matter deficits correlated with dyscalculia in children are only present early in development, and that over time these white matter deficits in dyscalculia become very small or even negligible.

In line with the only previous functional connectivity study in children with dyscalculia (Rosenberg-Lee et al., 2015), we found hyper-connectivity in adults with dyscalculia between FG and PVC, and between IOC and PVC. These regions are known to be involved in the processing of complex visual objects (Grill-Spector et al., 2008; Menon et al., 2000). It has been previously demonstrated that children with dyscalculia had decreased grey matter volume in these regions (Rykhlevskaia et al., 2009). Furthermore, the observed increased connectivity in this study was related to arithmetic skills, namely more functional connectivity is associated with lower arithmetic skills.

We suggest that these findings of increased functional connectivity might be interpreted in terms of compensatory processes that were activated during digit number processing in dyscalculia. More specifically, this increased functional connectivity from the occipital cortex to the infero-temporal cortex can be related to increased connectivity to the “visual number area” (located in the infero-temporal cortex). This area was first mentioned by Shum et al. (2013) who did intracranial electrophysiological recordings. Shum and colleagues (2013) demonstrated that this area responds more strongly to digits than to control conditions which are well-matched in terms of visual (letters, false fonts), semantic (number words) or phonological (phonologically similar non-number words) similarity. Furthermore, Srihasam et al., (2012) did a fMRI study in macaque monkeys and observed as well an area in the ventral temporal cortex selective for trained symbols (opposed to untrained shapes and faces). These results could suggest that such regions only develop when a high level of visual proficiency with number symbols is reached (as was the case for the younger monkeys in that study) (Piazza and Eger, 2016). Therefore, we suggest that the, in this study, observed increased connectivity in direction of the inferior-temporal cortex is caused by compensation mechanisms in DD to process Arabic digits. Note however that we dit not study the exact location of “visual number area”, as this region lies within or close to the fMRI signal-drop out zone produced by the nearby auditory canal and venous sinus artifacts Shum et al. (2013).

### 4.3. Anatomical differences

In line with previous studies in children (Rotzer et al., 2008; Rykhlevskaia et al., 2009), we found grey matter volume abnormalities for adults with dyscalculia in posterior cingulate cortex. The posterior cingulate cortex is known to be a part of the default mode network (DMN). The default mode network is typically deactivated during cognitive demanding tasks and assumed to be involved in efficiently processing external information and supporting mental activity that is internally directed (Raichle, 2015). Our findings in adults might be consistent with the interpretation of Rosenberg-Lee and colleagues (2015) that their findings are related to a possible deficit in the default mode network. However, important to note is that, against our expectations, we did not observe any anatomical abnormalities in the parietal cortex or the IPS in contrast to previous studies (Ranpura et al., 2013; Rotzer et al., 2008; Rykhlevskaia et al., 2009).

### 4.4. Domain-specific vs. domain-general deficits

Recently, there is an increasing awareness that the neural origins of dyscalculia are not restricted to domain-specific deficits such as impaired neural representations of number versus access deficit to these domain-relevant representations. Instead, or in addition, the neural correlates might also include domain-general deficits and involve impairments that are not specific to math) (Fias et al., 2013; Rubinsten and Henik, 2009; Wilson et al., 2015). For example, impairments in executive functions, working memory, attention or inhibitory control have been highlighted as potential risk factors for learning disorders and their comorbidity (Rubinsten and Henik, 2009).

Through our multi-faceted analyses, we were able to observe domain-specific as well as domain-general effects. We clearly observed domain-specific effects by showing impaired neural representations for non-symbolic magnitudes in adults with dyscalculia in brain regions typically correlated with numerical cognition (IPS and parietal cortex). On the other hand, we also observed domain-general effects. First of all, the impaired neural magnitude representations in dyscalculia were not *only* located in the IPS and parietal regions. As in previous studies, we demonstrated that deficits in functional neural correlates in dyscalculia are spread out throughout the cortex (Kaufmann et al., 2011; Rosenberg-Lee et al., 2015; Rotzer et al., 2008; Rykhlevskaia et al., 2009). In our study, the representations in (some of) these extra-parietal regions might be related to numerical representations as well as to how these representations are recruited during more domain-general processes. For example, frontal regions often play an important role in attention and working memory processes required for problem solving, also in numerical context. Secondly, our functional connectivity results showed hyper-connectivity for dyscalculia between visual regions and fusiform gyrus. Interestingly, increased functional connectivity was previously linked to compensation processes and inhibitory processes (Geerligs et al., 2012; Rosenberg-Lee et al., 2015). Thirdly, structural differences were located in the posterior cingulate cortex, a brain region that is part of the default mode network which is also linked to domain-general processes.

### 4.5. Conclusion

Our study supports a clear deficit in the quality of non-symbolic magnitude representations in adults with dyscalculia. On the other hand, we also observed hyper-connectivity in visual brain regions and increased grey matter volume in the default mode network in adults with dyscalculia. Hence, the deficits in dyscalculia cannot be localized to one brain region or to one particular type of brain deficit (functional, anatomical or connectivity). Our study illustrates the many advantages of combining different imaging techniques on a whole brain level to disentangle the etiology of neurodevelopmental disorders compared to focusing on only one brain region and/or only one neuroimaging method. We therefore suggest future studies to apply multi-method approaches to investigate neurodevelopmental disorders and mental disorders at large, in adults as well as children.

## 5. Acknowledgments

This work was supported by the Fund for Scientific Research Flanders (fellowship to J.B.), Wellcome Trust grant (to C.R.G. 098771/Z/12/Z and D.M. 101253/A/13/Z), an IDO Project of the KU Leuven (IDO/10/003), an FWO project (G.0946.12), a Federal Research Action (IUAP-P7/11), and an ERC grant (ERC-2011-Stg-284101). Recruitment of participants with dyscalculia was done with the help of the learning disorders diagnostic center PraxisP in Leuven, Belgium.

## References

American Psychiatric Association, 2013. Diagnostic and Statistical Manual of Mental Disorders. American Psychiatric Association. doi:10.1176/appi.books.9780890425596

Ansari, D., 2008. Effects of development and enculturation on number representation in the brain. Nat. Rev. Neurosci. 9, 278–291. doi:10.1038/nrn2334

Ashburner, J., Friston, K.J., 2000. Voxel-Based Morphometry—The Methods. Neuroimage 11, 805–821. doi:10.1006/nimg.2000.0582

Ashkenazi, S., Rosenberg-Lee, M., Tenison, C., Menon, V., 2012. Weak task-related modulation and stimulus representations during arithmetic problem solving in children with developmental dyscalculia. Dev. Cogn. Neurosci. 2 Suppl 1, S152–S166. doi:10.1016/j.dcn.2011.09.006

Balsters, J.H., Mantini, D., Apps, M.A.J., Eickhoff, S.B., Wenderoth, N., 2016. Connectivity-based parcellation increases network detection sensitivity in resting state fMRI: An investigation into the cingulate cortex in autism. NeuroImage Clin. 11, 494–507. doi:10.1016/j.nicl.2016.03.016

Baria, A.T., Mansour, A., Huang, L., Baliki, M.N., Cecchi, G.A., Mesulam, M.M., Apkarian, A.V., 2013. Linking human brain local activity fluctuations to structural and functional network architectures. Neuroimage 73, 144–155. doi:10.1016/j.neuroimage.2013.01.072

Bishop, D.V.M., 2010. Which Neurodevelopmental Disorders Get Researched and Why? PLoS One 5, e15112. doi:10.1371/journal.pone.0015112

Boets, B., Op de Beeck, H.P., Vandermosten, M., Scott, S.K., Gillebert, C.R., Mantini, D., Bulthe, J., Sunaert, S., Wouters, J., Ghesquiere, P., 2013. Intact But Less Accessible Phonetic Representations in Adults with Dyslexia. Science (80-.). 342, 1251–1254. doi:10.1126/science.1244333

Brainard, D.H., 1997. The Psychophysics Toolbox. Spat. Vis. 10, 433–436. doi:10.1163/156856897X00357

Brus, B., 1999. Een-minuut-test.

Bulthé, J., De Smedt, B., Op de Beeck, H.P., 2015. Visual Number Beats Abstract Numerical Magnitude: Format-dependent Representation of Arabic Digits and Dot Patterns in Human Parietal Cortex. J. Cogn. Neurosci. 27, 1376–1387. doi:10.1162/jocn_a_00787

Bulthé, J., De Smedt, B., Op de Beeck, H.P., 2014a. Format-dependent representations of symbolic and non-symbolic numbers in the human cortex as revealed by multi-voxel pattern analyses. Neuroimage 87, 311–322. doi:10.1016/j.neuroimage.2013.10.049

Bulthé, J., van den Hurk, J., Daniels, N., De Smedt, B., Op de Beeck, H.P., 2014b. A validation of a multi-spatialscale method for multivariate pattern analysis, in: 2014 International Workshop on Pattern Recognition in Neuroimaging. IEEE, pp. 1–4. doi:10.1109/PRNI.2014.6858513

Butterworth, B., Varma, S., Laurillard, D., 2011. Dyscalculia: From Brain to Education. Science (80-.). 332, 1049–1053. doi:10.1126/science.1201536

Chang, C.-C., Lin, C.-J., 2011. {LIBSVM}: A library for support vector machines. ACM Trans. Intell. Syst. Technol. 2, 27:1--27:27. doi:http://doi.acm.org/10.1145/1961189.1961199

Cohen Kadosh, R., Cohen Kadosh, K., Kaas, A., Henik, A., Goebel, R., 2007. Notation-dependent and -independent representations of numbers in the parietal lobes. Neuron 53, 307–14. doi:10.1016/j.neuron.2006.12.025

Damarla, S.R., Just, M.A., 2013. Decoding the representation of numerical values from brain activation patterns. Hum. Brain Mapp. 34, 2624–2634. doi:10.1002/hbm.22087

De Smedt, B., Boets, B., 2010. Phonological processing and arithmetic fact retrieval: Evidence from developmental dyslexia. Neuropsychologia 48, 3973–3981. doi:10.1016/j.neuropsychologia.2010.10.018

De Smedt, B., Gilmore, C.K., 2011. Defective number module or impaired access? Numerical magnitude processing in first graders with mathematical difficulties. J. Exp. Child Psychol. 108, 278–292. doi:10.1016/j.jecp.2010.09.003

De Smedt, B., Noël, M.-P., Gilmore, C., Ansari, D., 2013. How do symbolic and non-symbolic numerical magnitude processing skills relate to individual differences in children’s mathematical skills? A review of evidence from brain and behavior. Trends Neurosci. Educ. 2, 48–55. doi:10.1016/j.tine.2013.06.001

De Vos, T., 1992. Tempo-Test-Rekenen. Handleiding [Tempo Test Arithmetic. Manual]. Nijmegen: Berkhout.

Dehaene, S., Izard, V., Piazza, M., 2005. Control over non-numerical parameters in numerosity experiments. Unpubl. Manuscr. (available www.unicog.org).

Ebisch, S.J.H., Mantini, D., Romanelli, R., Tommasi, M., Perrucci, M.G., Romani, G.L., Colom, R., Saggino, A., 2013. Long-range functional interactions of anterior insula and medial frontal cortex are differently modulated by visuospatial and inductive reasoning tasks. Neuroimage 78, 426–438. doi:10.1016/j.neuroimage.2013.04.058

Eger, E., Michel, V., Thirion, B., Amadon, A., Dehaene, S., Kleinschmidt, A., 2009. Deciphering cortical number coding from human brain activity patterns. Curr. Biol. 19, 1608–1615. doi:10.1016/j.cub.2009.08.047

Elsabbagh, M., Divan, G., Koh, Y.-J., Kim, Y.S., Kauchali, S., Marcín, C., Montiel-Nava, C., Patel, V., Paula, C.S., Wang, C., Yasamy, M.T., Fombonne, E., 2012. Global Prevalence of Autism and Other Pervasive Developmental Disorders. Autism Res. 5, 160–179. doi:10.1002/aur.239

Estrada-Mejia, C., de Vries, M., Zeelenberg, M., 2016. Numeracy and wealth. J. Econ. Psychol. 54, 53–63. doi:10.1016/j.joep.2016.02.011

Feigenson, L., Dehaene, S., Spelke, E., 2004. Core systems of number. Trends Cogn. Sci. 8, 307–314. doi:10.1016/j.tics.2004.05.002

Fias, W., Menon, V., Szucs, D., 2013. Multiple components of developmental dyscalculia. Trends Neurosci. Educ. 2, 43–47. doi:10.1016/j.tine.2013.06.006

French, J., Ekstrom, R., Price, L., 1963. Manual for kit of reference tests for cognitive factors (revised 1963) (Tech. Rep.). DTIC Document.

Geerligs, L., Saliasi, E., Maurits, N.M., Lorist, M.M., 2012. Compensation through Increased Functional Connectivity: Neural Correlates of Inhibition in Old and Young. J. Cogn. Neurosci. 24, 2057–2069. doi:10.1162/jocn_a_00270

Gerardi, K., Goette, L., Meier, S., 2013. Numerical ability predicts mortgage default. Proc. Natl. Acad. Sci. 1–5. doi:10.1073/pnas.1220568110

Good, C.D., Johnsrude, I.S., Ashburner, J., Henson, R.N., Friston, K.J., Frackowiak, R.S., 2001. A voxel-based morphometric study of ageing in 465 normal adult human brains. Neuroimage 14, 21–36. doi:10.1006/nimg.2001.0786

Grill-Spector, K., Golarai, G., Gabrieli, J., 2008. Developmental neuroimaging of the human ventral visual cortex. Trends Cogn. Sci. 12, 152–162. doi:10.1016/j.tics.2008.01.009

Hebart, M.N., Görgen, K., Haynes, J.-D., Dubois, J., 2015. The Decoding Toolbox (TDT): a versatile software package for multivariate analyses of functional imaging data. Front. Neuroinform. 8, 1–18. doi:10.3389/fninf.2014.00088

Holloway, I.D., Battista, C., Vogel, S.E., Ansari, D., 2013. Semantic and Perceptual Processing of Number Symbols: Evidence from a Cross-linguistic fMRI Adaptation Study. J. Cogn. Neurosci. 25, 388–400. doi:10.1162/jocn_a_00323

Johnson, M.H., Halit, H., Grice, S.J., Karmiloff-Smith, A., 2002. Neuroimaging of typical and atypical development: a perspective from multiple levels of analysis. Dev. Psychopathol. 14, 521–536. doi:10.1017/S0954579402003073

Kaufmann, L., Wood, G., Rubinsten, O., Henik, A., 2011. Meta-Analyses of Developmental fMRI Studies Investigating Typical and Atypical Trajectories of Number Processing and Calculation. Dev. Neuropsychol. 36, 763–787. doi:10.1080/87565641.2010.549884

Kriegeskorte, N., Bandettini, P., 2007. Analyzing for information, not activation, to exploit high-resolution fMRI. Neuroimage 38, 649–662. doi:10.1016/j.neuroimage.2007.02.022

Kriegeskorte, N., Goebel, R., Bandettini, P., 2006. Information-based functional brain mapping. PNAS 103, 3863–3868. doi:10.1073/pnas.0600244103

Kucian, K., Ashkenazi, S.S., Hänggi, J., Rotzer, S., Jäncke, L., Martin, E., von Aster, M., 2013. Developmental dyscalculia: a dysconnection syndrome? Brain Struct. Funct. doi:10.1007/s00429-013-0597-4

Leemans, A., Jeurissen, B., Sijbers, J., Jones, D.K., 2009. ExploreDTI: a graphical toolbox for processing, analyzing, and visualizing diffusion MR data, in: 17th Annual Meeting of Intl Soc Mag Reson Med. Hawaii, USA, p. 3537.

Lyons, I.M., Ansari, D., 2009. The cerebral basis of mapping nonsymbolic numerical quantities onto abstract symbols: an fMRI training study. J. Cogn. Neurosci. 21, 1720–35. doi:10.1162/jocn.2009.21124

Lyons, I.M., Ansari, D., Beilock, S.L., 2015. Qualitatively different coding of symbolic and nonsymbolic numbers in the human brain. Hum. Brain Mapp. 36, 475–488. doi:10.1002/hbm.22641

Matejko, A. a., Ansari, D., 2014. Drawing connections between white matter and numerical and mathematical cognition: A literature review. Neurosci. Biobehav. Rev. 48, 35–52. doi:10.1016/j.neubiorev.2014.11.006

Menon, V., 2011. Large-scale brain networks and psychopathology: A unifying triple network model. Trends Cogn. Sci. 15, 483–506. doi:10.1016/j.tics.2011.08.003

Menon, V., Rivera, S.M.M., White, C.D.D., Glover, G.H.H., Reiss, A.L.L., 2000. Dissociating Prefrontal and Parietal Cortex Activation during Arithmetic Processing. Neuroimage 12, 357–365. doi:10.1006/nimg.2000.0613

Misaki, M., Kim, Y., Bandettini, P. a, Kriegeskorte, N., 2010. Comparison of multivariate classifiers and response normalizations for pattern-information fMRI. Neuroimage 53, 103–118. doi:10.1016/j.neuroimage.2010.05.051

Mourão-Miranda, J., Bokde, A.L.W., Born, C., Hampel, H., Stetter, M., 2005. Classifying brain states and determining the discriminating activation patterns: Support Vector Machine on functional MRI data. Neuroimage 28, 980–995. doi:10.1016/j.neuroimage.2005.06.070

Mussolin, C., De Volder, A., Grandin, C., Schlögel, X., Nassogne, M.-C., Noël, M.-P., 2010a. Neural Correlates of Symbolic Number Comparison in Developmental Dyscalculia. J. Cogn. Neurosci. 22, 860–874. doi:10.1162/jocn.2009.21237

Mussolin, C., Mejias, S., Noël, M.-P., 2010b. Symbolic and nonsymbolic number comparison in children with and without dyscalculia. Cognition 115, 10–25. doi:10.1016/j.cognition.2009.10.006

Noël, M.-P., Rousselle, L., 2011. Developmental Changes in the Profiles of Dyscalculia: An Explanation Based on a Double Exact-and-Approximate Number Representation Model. Front. Hum. Neurosci. 5, 1–4. doi:10.3389/fnhum.2011.00165

Piazza, M., Eger, E., 2016. Neural foundations and functional specificity of number representations. Neuropsychologia 83, 257–273. doi:10.1016/j.neuropsychologia.2015.09.025

Piazza, M., Izard, V., Pinel, P., Le Bihan, D., Dehaene, S., 2004. Tuning curves for approximate numerosity in the human intraparietal sulcus. Neuron 44, 547–555. doi:10.1016/j.neuron.2004.10.014

Piazza, M., Pinel, P., Le Bihan, D., Dehaene, S., 2007. A magnitude code common to numerosities and number symbols in human intraparietal cortex. Neuron 53, 293–305. doi:10.1016/j.neuron.2006.11.022

Pinel, P., Piazza, M., Le Bihan, D., Dehaene, S., 2004. Distributed and overlapping cerebral representations of number, size, and luminance during comparative judgments. Neuron 41, 983–993.

Power, J.D., Barnes, K.A., Snyder, A.Z., Schlaggar, B.L., Petersen, S.E., 2012. Spurious but systematic correlations in functional connectivity MRI networks arise from subject motion. Neuroimage 59, 2142–2154. doi:10.1016/j.neuroimage.2011.10.018

Price, G.R., Holloway, I., Rä sänen, P., Vesterinen, M., Ansari, D., 2007. Impaired parietal magnitude processing in developmental dyscalculia. Curr. Biol. 17, R1042–R1043. doi:10.1016/j.cub.2007.10.013

Raichle, M.E., 2015. The Brain’s Default Mode Network. Annu. Rev. Neurosci. 38, 433–447. doi:10.1146/annurev-neuro-071013-014030

Ranpura, A., Isaacs, E., Edmonds, C., Rogers, M., Lanigan, J., Singhal, A., Clayden, J., Clark, C., Butterworth, B., 2013. Developmental trajectories of grey and white matter in dyscalculia. Trends Neurosci. Educ. 2, 56–64. doi:10.1016/j.tine.2013.06.007

Reyna, V.F., Nelson, W.L., Han, P.K., Dieckmann, N.F., 2009. How numeracy influences risk comprehension and medical decision making. Psychol. Bull. 135, 943–73. doi:10.1037/a0017327

Ritchie, S.J., Bates, T.C., 2013. Enduring Links From Childhood Mathematics and Reading Achievement to Adult Socioeconomic Status. Psychol. Sci. 24, 1301–1308. doi:10.1177/0956797612466268

Rosenberg-Lee, M., Ashkenazi, S., Chen, T., Young, C.B., Geary, D.C., Menon, V., 2015. Brain hyper-connectivity and operation-specific deficits during arithmetic problem solving in children with developmental dyscalculia. Dev. Sci. 18, 351–372. doi:10.1111/desc.12216

Rotzer, S., Kucian, K., Martin, E., Aster, M. von, Klaver, P., Loenneker, T., 2008. Optimized voxel-based morphometry in children with developmental dyscalculia. Neuroimage 39, 417–422. doi:10.1016/j.neuroimage.2007.08.045

Rousselle, L., Noël, M.-P., 2007. Basic numerical skills in children with mathematics learning disabilities: a comparison of symbolic vs non-symbolic number magnitude processing. Cognition 102, 361–395. doi:10.1016/j.cognition.2006.01.005

Rubinsten, O., Henik, A., 2009. Developmental dyscalculia: heterogeneity might not mean different mechanisms. Trends Cogn. Sci. 13, 92–9. doi:10.1016/j.tics.2008.11.002

Rykhlevskaia, E., Uddin, L.Q., Kondos, L., Menon, V., 2009. Neuroanatomical correlates of developmental dyscalculia: combined evidence from morphometry and tractography. Front. Hum. Neurosci. 3, 1–13. doi:10.3389/neuro.09.051.2009

Shum, J., Hermes, D., Foster, B.L., Dastjerdi, M., Rangarajan, V., Winawer, J., Miller, K.J., Parvizi, J., 2013. A brain area for visual numerals. J. Neurosci. 33, 6709–6715. doi:10.1523/JNEUROSCI.4558-12.2013

Simos, P.G., Kanatsouli, K., Fletcher, J.M., Sarkari, S., Juranek, J., Cirino, P., Passaro, A., Papanicolaou, A.C., 2008. Aberrant spatiotemporal activation profiles associated with math difficulties in children: A magnetic source imaging study. Neuropsychology 22, 571–584. doi:10.1037/0894-4105.22.5.571

Spreen, O., Risser, A., Edgell, D., 1995. Developmental Neuropsychology. Oxford University Press, New York.

Srihasam, K., Mandeville, J.B., Morocz, I. a, Sullivan, K.J., Livingstone, M.S., 2012. Behavioral and anatomical consequences of early versus late symbol training in macaques. Neuron 73, 608–619. doi:10.1016/j.neuron.2011.12.022

Ung, H., Brown, J.E., Johnson, K.A., Younger, J., Hush, J., Mackey, S., 2014. Multivariate classification of structural MRI data detects chronic low back pain. Cereb. Cortex 24, 1037–1044. doi:10.1093/cercor/bhs378

Van den Bos, K., 1999. De Klepel.

Wang, R., Wedeen, V.J., 2007. TrackVis.org, Martinos Center for Biomedical Imaging, Massachusetts General Hospital., in: Proc Int Soc Magn Reson Med 15. p. 3720.

Wilson, A.J., Andrewes, S.G., Struthers, H., Rowe, V.M., Bogdanovic, R., Waldie, K.E., 2015. Dyscalculia and dyslexia in adults: Cognitive bases of comorbidity. Learn. Individ. Differ. 37, 118–132. doi:10.1016/j.lindif.2014.11.017

Xia, M., Wang, J., He, Y., 2013. BrainNet Viewer: a network visualization tool for human brain connectomics. PLoS One 8, e68910. doi:10.1371/journal.pone.0068910

Zorzi, M., Di Bono, M.G., Fias, W., 2011. Distinct representations of numerical and non-numerical order in the human intraparietal sulcus revealed by multivariate pattern recognition. Neuroimage 56, 674–680. doi:10.1016/j.neuroimage.2010.06.035

